# HDL-C as a potential medium between depletion of *Lachnospiraceae* genera and hypertension under high-calorie diet

**DOI:** 10.1101/2022.06.21.497117

**Authors:** Yongmei Lan, Kang Ning, Yanqing Ma, Jin Zhao, Caihong Ci, Xiao Yang, Fulong An, Zilong Zhang, Yan An, Mingyue Cheng

## Abstract

Gut microbial dysbiosis has been associated with hypertension. An extremely high incidence of essential hypertension was found in the Han and the Yugur who resided in Sunan county in East Asia’s nomadic steppes with little population movement. In attempt to investigate the gut microbial role in hypertension, we recruited a total of 1, 242 Yugur and Han people, who had resided in Sunan County for more than 15 years and accounted for 3% of the local population. The epidemiological survey of 1,089 individuals indicated their nearly 1.8 times higher prevalence of hypertension (38.2–43.3%) than the average in China (23.2%), under a special high-calorie diet based on wheat, cattle, mutton, and animal offal. The 16S rRNA gene sequencing on the fecal samples of 153 individuals revealed that certain *Lachnospiraceae* genera were negatively correlated with high-density lipoprotein cholesterol (HDL-C, *P* = 5.46 × 10^−6^), systolic blood pressure (SBP, *P* = 7.22 × 10^−3^), diastolic blood pressure (DBP, *P* = 1.8 × 10^−3^). HDL-C was positively correlated with SBP (*P* = 0.023). We further observed that serum butyrate content was lower in both Han (*P* = 1.99 × 10^−3^) and Yugur people (*P* = 0.031) with hypertension than those without hypertension. This study gives a novel insight into the role of gut microbial dysbiosis in hypertension modulation under a high-calorie diet, where the notable depletion of *Lachnospiraceae* genera might lead to less production of butyrate, contributing to the lower level of HDL-C, and elevating blood pressure in hypertension.

**IMPORTANCE:** Dietary nutrients can be converted by gut microbiota into metabolites such as short-chain fatty acids, which may serve as disease-preventing agents in hypertension. Due to limited population mobility and a unique high-calorie dietary habit, the recruited cohort in this study could be a representative for elucidating the associations between gut microbiota and hypertension under high-calorie diet. Moreover, low levels of HDL-C have previously been associated with an increased risk of various cardiovascular diseases (CVDs). Our findings provide a new insight that low levels of HDL-C may be a potential medium between depletion of *Lachnospiraceae* genera and hypertension under high-calorie diet, which might also be a potential candidate for other CVDs.

## INTRODUCTION

Human gut microbiota has been correlated with a variety of cardiovascular diseases (CVDs) pathogenesis, such as hypertension (1, 2). Hypertension is a major modifiable risk factor for the CVD such as myocardial infarction, heart failure, and stroke (3, 4). Compared to genetic effects that contribute less than 20% to the risk of developing CVD pathogenesis, environmental effects especially diet are known as the prominent role in CVD pathogenesis (1, 5–7). Additionally, a diet rich in fruits, vegetables, and low-fat dairy products and reduced saturated and total fat has been confirmed to ameliorate hypertension in multiple randomized controlled trials (8). Moreover, gut microbiota, whose composition are dominantly modulated by diet (9, 10), can convert dietary nutrients into metabolites such as short-chain fatty acids (SCFA) that acts as the potential disease-preventing factors in hypertension (1, 11). Indeed, our epidemiology survey showed that local Yugur and Han people, who resided in Sunan county in East Asia’s nomadic steppes with little population movement, followed high-calorie dietary custom and presented extremely higher incidence of essential hypertension (**Table S1**). Here, we have investigated gut microbiota of local Han and Yugur people, with or without essential hypertension, to gain insight into the potential microbial contribution to their high incidence of hypertension.

The Yugur, one of East Asian ethnic groups with a population of only 14,378, emerged around the eighth century by gathering mainly the Hexi Uighur and a few Mongolian, Tibetan and other ethnics. The Yugur reside in Sunan County that is located in the middle of Hexi Corridor, at the north foot of Qilian Mountain in Northwest China, with a length of more than 650 kilometers and an average altitude of 3,200 meters (**Fig. 1A**). This area has an alpine semi-arid climate with an annual average temperature of 4 ℃, and the terrain is relatively closed and sparsely populated. Due to the unique natural environment, the Yugur have developed a special high-calorie diet based on wheat, cattle, mutton, animal offal, dairy products, and Chinese Baijiu with limited intake of vegetables and fruits. Moreover, since the establishment of Sunan County in 1954, the Han who have successively immigrated to this area have been assimilated to Yugur’s customs, sharing a similar high-calorie diet. The high-calorie diet might be one of the major causes of their high incidence of hypertension. According to our recent epidemiological survey of essential hypertension in Sunan county, the prevalence of essential hypertension among Yugurs was 43.3% and among Hans was 38.2%, both of which were higher than China’s national average (23.2%, 2012–2015) (12). The high-calorie diet may also equip Yugur and Han individuals with a distinct gut microbial composition, therefore influencing the pathogenesis of hypertension, but the gut microbial patterns and regulatory mechanisms behind this proposed modulating process remain unknown.

**FIG 1.**
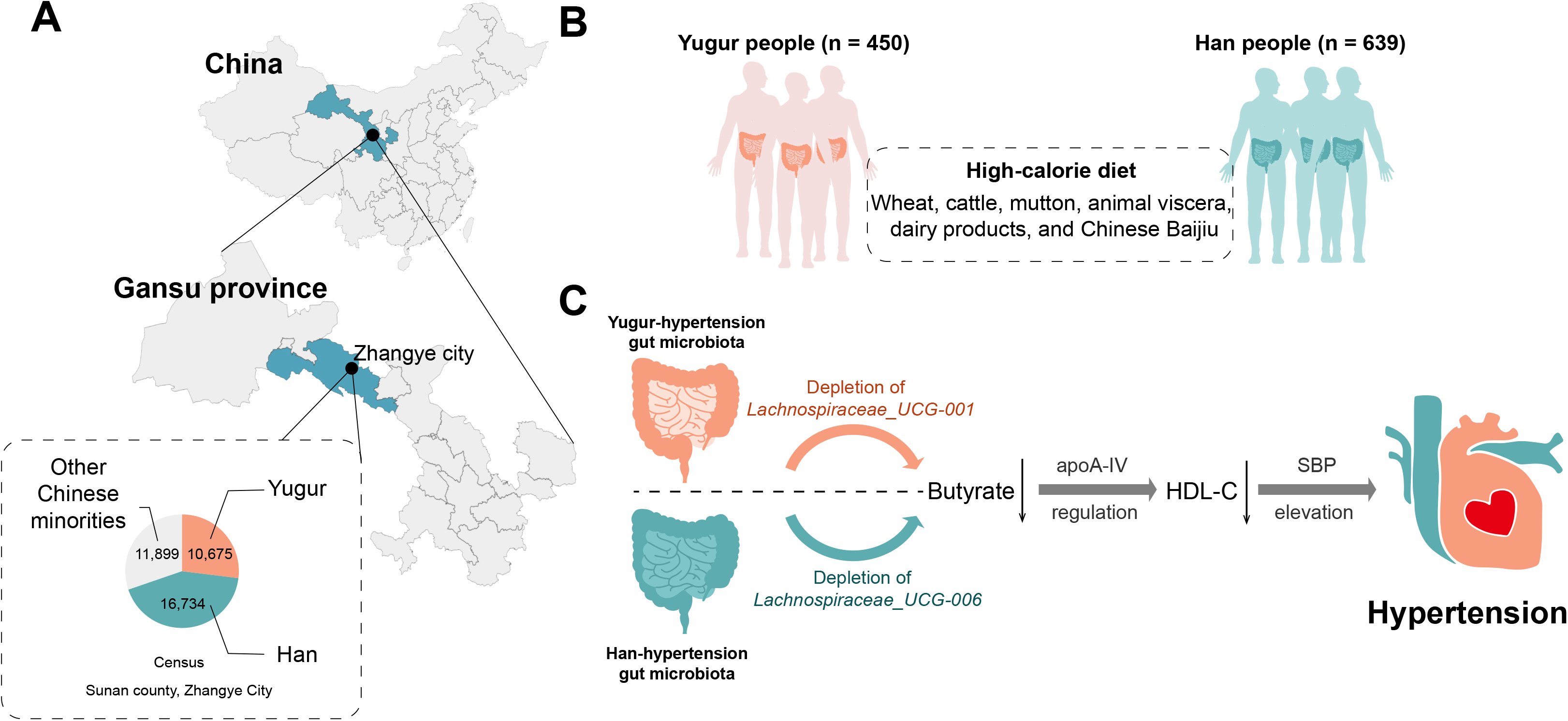
The recruited cohort and the potential link between gut microbiota and hypertension. (A) The recruited cohort resided in Sunan County, Gansu Province, China. The local populational proportion is shown in the pie chart. (B) Yugur people (n = 450) and Han people (n = 639), who had been living in Sunan County for more than 15 years, were investigated for the epidemic survey of hypertension prevalence and dietary custom in this study. (C) The potential ethnic-specific mechanism proposed in this study that gut microbiota might promote hypertension pathogenesis by reducing intestinal butyrate and lowering HDL-C.

In this study, we investigated a total of 1, 242 Yugur and Han people who had lived in Sunan County for more than 15 years and accounted for 3% of the local population, to investigate the association and possible mechanism of gut microbiota in the pathogenesis of hypertension under high-calorie diet. Due to limited population mobility and a unique high-calorie dietary habit, this cohort could be representative for elucidating the associations between gut microbiota and hypertension in the presence of a high-calorie diet.

## RESULTS

### Han and Yugur people in Sunan County present higher prevalence of hypertension with high-calorie diet

We conducted an epidemic survey of essential hypertension and investigated the dietary structure on a total of 1,089 individuals in Sunan County in 2015, including 639 Han people and 450 Yugur people (**Table S1, Fig. 1B**). The prevalence of hypertension was 38.2% in Han people, which was lower than that of Yugur people with 43.3%. Moreover, compared to the Dietary Guidelines for Chinese Residents (13), both Han and Yugur shared a high-calorie diet: 1) Excessive intake of meat (178.8 ∼ 234.9g/d) that was more than the requirement of the Chinese dietary guidelines (50 ∼ 100g/d); and 2) Limited intake of vegetables and fruits (325.7 ∼ 387.5g/d) that was less than the requirement of the Chinese dietary guidelines (500 ∼ 700g/d). Furthermore, compared to people without hypertension, people with hypertension consumed more beef and mutton, animal offal, fried food, milk and its products, edible oil, and Chinese Baijiu. The high-calorie diet might play a crucial role in higher prevalence of hypertension in Han and Yugur people in Sunan County.

### Gut microbiota was of dysbiosis in Han and Yugur people with hypertension

We then collected 153 fecal samples of Han and Yugur people who has been living in Sunan County for at least 15 years to examine their gut microbial compositions. We found several dietary factors were correlated with their microbial compositions (*P* < 0.05, PERMANOVA, **Table S2**), such as wheat, rice, coarse cereals, vegetable and fruits, animal offal, butter, and edible oil. To explore differences in microbial composition between hypertension and non-hypertension, we firstly performed principal coordinate analysis (PCoA) on all of fecal samples using unweighted (**Fig. 2A**) and weighted Unifrac distance (**Fig. 2B**). We found that hypertension samples were evidently separated from non-hypertension samples against PCo1 axis when both distances were used (*P* = 5.08 × 10^−5^, *P* = 4.48 × 10^−3^). In addition, Yugur people without hypertension had higher microbial Shannon diversity than that of Han people (*P* = 0.016), though microbial Shannon diversity showed no significant difference between hypertension and non-hypertension in both ethnic groups (**Fig. 2C**).

**FIG 2.**
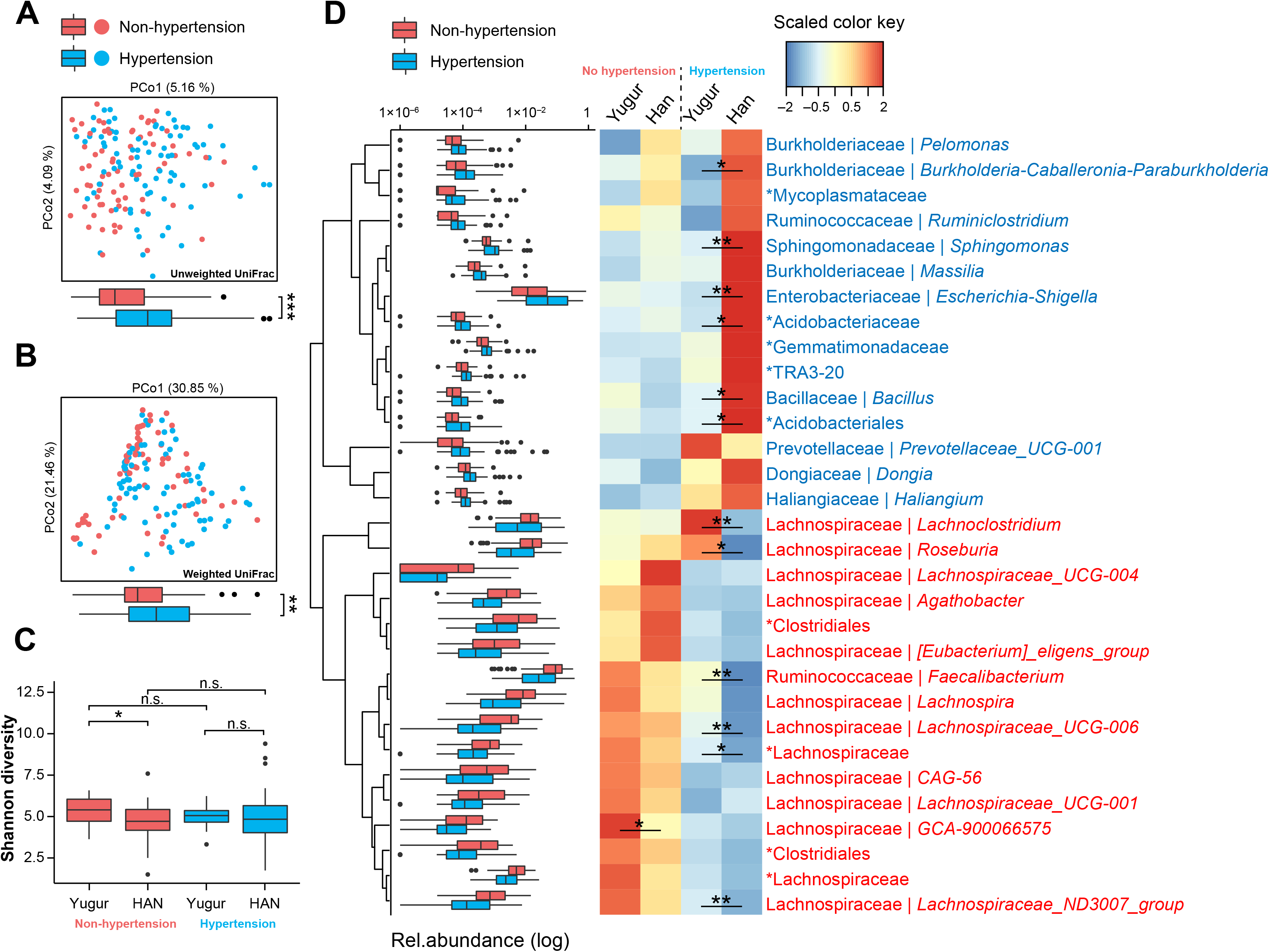
Differences in microbiota composition between hypertension and non-hypertension individuals from Han and Yugur. (A) and (B) Individual gut microbiota compositions of 81 hypertension patients and 72 non-hypertension individuals plotted on an unweighted UniFrac PCoA plot (A) and a weighted UniFrac PCoA plot (B), with a boxplot below each one showing sample distributions. (C) The boxplot shows microbiota Shannon diversity of 17 Yugur without hypertension, 55 Han without hypertension, 23 Yugur with hypertension, and 58 Han with hypertension. (D) Boxplots in the left panel show relative abundances of 31 specific genera that had significantly different distributions between hypertension and non-hypertension groups (*P* < 0.05, q < 0.05, Mann-Whitney-Wilcoxon test). Hierarchical Ward-linkage clustering was based on the Euclidean distance of these genera’s abundance among all the 153 samples. The heatmap in the right panel showed the scaled mean abundances of these genera in four subgroups as described in (C). The significance between subgroups were annotated inner the heatmap. The classified genera were annotated with the family and genus name, and the unclassified genera were designated as a higher rank with an asterisk. In all the panels, statistical significance was tested using the Mann-Whitney-Wilcoxon test (*, *P* < 0.05; **, *P* < 0.01). The boxes represent 25th–75th percentiles, black lines indicate the median and whiskers extend to the maximum and minimum values within 1.5 × the interquartile range.

We then identified a total of 5 microbial phyla, 8 classes, 23 orders, 36 families, 54 genera that were significantly elevated or depleted (P < 0.05, q < 0.1, Mann-Whitney-Wilcoxon test) in gut microbiota of people with hypertension (shortened as hypertension microbiota), as compared to that of people without hypertension (shortened as non-hypertension microbiota) (**Fig. 2D, Table S3**). Among these 54 genera, 31 genera with q < 0.05 were designated as hypertension-related genera. A total of 15 hypertension-related genera were found significantly elevated in hypertension microbiota, such as *Ruminiclostridium* (*P* = 4.56 × 10^−3^, q = 0.036) whose metabolic pathways were related to blood pressure regulation (14), and *Escherichia-Shigella* (*P* = 3.20 × 10^−4^, q = 4.14 × 10^−3^) whose infection in gastroenteritis was correlated with an increased risk of hypertension (15). Moreover, we observed elevation of *Pelomonas* (*P* = 1.63 × 10^−3^, q = 0.015) and *Sphingomonas* (*P* = 1.86 × 10^−4^, q = 0.038) that have been reported to be found in blood microbiome and positively correlated with a few inflammatory markers (16), and the risk of hypertension (17), respectively. It was speculated that these two gut microbes might be transited to blood microbiome to promote hypertension, under circumstance of increased gut permeability in people with hypertension (18), which deserved further investigations.

### Depletion of Lachnospiraceae genera dominates microbial dysbiosis in Han and Yugur people with hypertension

Notably, a total of 16 hypertension-related genera, significantly depleted in hypertension microbiota, were found mostly from the family Lachnospiraceae (**Fig. 2D, Table S3**), such as *Lachnospiraceae_UCG-001* (*P* = 6.50 × 10^−3^, q = 0.046), *Lachnospiraceae_UCG-004* (*P* = 1.48 × 10^−4^, q = 3.77 × 10^− 3^), *Lachnospiraceae_UCG-006* (*P* = 9.37 × 10^−8^, q = 2.06 × 10^−5^), *Lachnospira* (*P* = 1.18 × 10^−5^, 8.67 × 10^−4^), *Agathobacter* (*P* = 2.74 × 10^−5^, q = 1.50 × 10^−3^), *Faecalibacterium* (*P* = 1.41 × 10^−4^, q = 3.77 × 10^−3^), and *Roseburia* (*P* = 2.18 × 10^−4^, q = 4.00 × 10^−3^). Gut microbes belonging to the family Lachnospiraceae were reported to impact human hosts by producing short-chain fatty acids, converting primary to secondary bile acids (19–21), and facilitating colonization resistance against intestinal pathogens (22, 23). *Roseburia* species, for instance, have been reported to protect against atherosclerosis by generating butyrate (24). These results implied a crucial role of Lachnospiraceae genera in pathogeneisis of hypertension in Han and Yugur people.

### Yugur people with hypertension presented less altered microbiota

We noticed that among the 31 hypertension-related genera identified in our study, only significant elevation of *Haliangium* (*P* = 0.042), and only significant depletion of *Lachnospiraceae_UCG-001* (*P* = 4.43 × 10^−3^), *GCA-900066575* (*P* = 3.31 × 10^−3^), and two unclassified genera of family Lachnospiraceae (*P* = 0.034) and order Clostridiales (*P* = 0.032), respectively, were observed in hypertension microbiota compared to non-hypertension microbiota, when investigating only Yugur people (**Table S4**). Nevertheless, except *Lachnospiraceae_UCG-001* (*P* = 0.18), all of other 30 genera kept significant elevation or depletion when investigating only Han people (**Table S4**).

Moreover, a certain number of genera in Yugur-hypertension microbiota, were found less altered than those in Han-hypertension microbiota (**Fig. 2D**). Compared to Han-hypertension microbiota, *Burkholderia-Caballeronia-Paraburkholderia* (*P* = 0.017), *Sphingomonas* (*P* = 8.97 × 10^−3^), *Escherichia-Shigella* (*P* = 2.51 × 10^−3^), *Bacillus* (*P* = 0.021), and two unclassified genera of order Acidobacteriales (*P* = 0.036, *P* = 0.018) were less elevated in Yugur-hypertension microbiota. In addition, *Lachnoclostridium* (*P* = 2.19 × 10^−3^, *Roseburia* (*P* = 0.032), *Faecalibacterium* (*P* = 5.08 × 10^−3^), *Lachnospiraceae_UCG-006* (*P* = 3.09 × 10^−3^), *Lachnospiraceae_ND3007_group* (*P* = 4.04 × 10^−3^), and an unclassified genus of family Lachnospiraceae (*P* = 0.035) were less depleted in Yugur-hypertension microbiota. These results suggested that Yugur people with hypertension had less altered microbiota, though the statistical significance might be biased by the different cohort size.

### Most discriminant microbial features of Han and Yugur people with hypertension

To explore the most discriminant microbes between hypertension and non-hypertension groups, we then performed random forest algorithm on the whole cohort (n = 153), Han people (n = 113), Yugur people (n = 40), respectively (**Fig. 3A**). Han-hypertension microbiota could be discriminated from Han-non-hypertension microbiota with the best AUROC (0.7884), while the AUROC was only 0.6522 when applying to Yugur people. Lachnospiraceae genera were the most discriminant features for each ethnic group to discriminate hypertension microbiota from non-hypertension microbiota (**Fig. 3B**, **Table S5–7**), with *Lachnospiraceae_UCG-006* for Han group and *Lachnospiraceae_UCG-001* for Yugur group. Moreover, we found family Lachnospiraceae was largely depleted in Han-hypertension microbiota (*P* = 1.6 × 10^−3^), while maintained the same level in Yugur-hypertension microbiota as it in non-hypertension microbiota (**Fig. 3C**). Moreover, *Lachnospiraceae_UCG-006* was notably depleted in Han-hypertension microbiota (*P* = 1.7 × 10^−8^, **Fig. 3D**) but not in Yugur-hypertension microbiota. On the contrary, *Lachnospiraceae_UCG-001* was significantly depleted in Yugur-hypertension microbiota (*P* = 4.4 × 10^−3^, **Fig. 3E**) but not in Han-hypertension microbiota.

**FIG 3.**
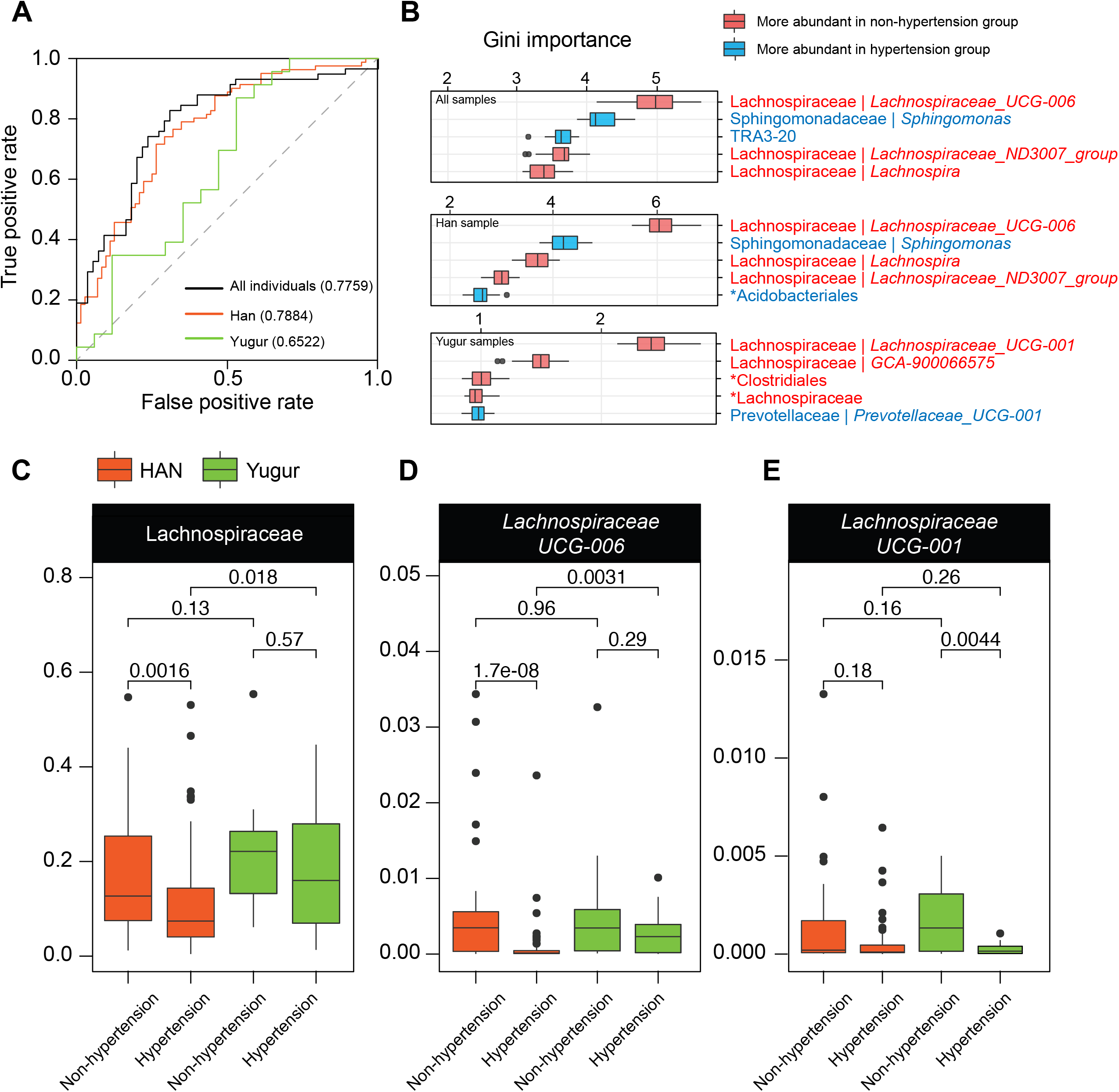
Microbial biomarkers for discriminating hypertension from non-hypertension. Randomforest algorithm with 10 randomized 10-fold cross validation was performed on 31 hypertension-related genera identified in Fig. 2, using all samples (n = 153), HAN samples (n = 113), Yugur samples (n = 40), to calculate Area Under the Receiver Operating Characteristic curve (AUROC), respectively (A), and Gini importance of each genus feature (B). The top five features are displayed and colored to show which group they are more significantly abundant in (*P* < 0.05, q < 0.05, Mann-Whitney-Wilcoxon test). (C) Abundances of the most discriminant genus features and their family among hypertension and non-hypertension groups of HAN and Yugur are plotted using boxplots. Statistical significance was calculated by Mann-Whitney-Wilcoxon test. Boxes represent the interquartile range between first and third quartiles and the line inside represents the median. Whiskers denote the lowest and highest values within 1.5 × interquartile range from the first and third quartiles, respectively.

### Depletion of Lachnospiraceae genera might promote hypertension by lowering the serum level of HDL-C

We subsequently explore the correlations of the recognized hypertension-related microbes with people physiological properties (**Fig. 4**). Nine physiological properties were found significantly changed in people with hypertension, compared to people without hypertension (*P* < 0.05, q < 0.1, Mann-Whitney-Wilcoxon test, **Table S8, Fig. 4A**). We performed Spearman correlation analysis on these nine physiological properties with 31 hypertension-related genera and 20 hypertension-related families, respectively (**Table S3** and **Table S9**). A total of eight properties were significantly correlated with 30 microbial taxa (*P* < 0.05, q < 0.1, Spearman correlation, **Table S9**). Systolic blood pressure (SBP) was positively correlated with family Mycoplasmataceae (r = 0.24, *P* = 4.43 × 10^−3^, q = 0.089) and genus *Escherichia-Shigella* (r = 0.22, *P* = 8.49 × 10^−3^, q = 0.053), while negatively correlated with genus *Lachnospiraceae_UCG-006* (r = –0.22, *P* = 8.27 × 10^−3^, q = 0.052) and *Lachnospiraceae _ND3007_group* (r = – 0.22, *P* = 7.22 × 10^−3^, q = 0.051). Diastolic blood pressure (DBP) was negatively correlated with four genera of family Lachnospiraceae including *GCA-900066575* (r = – 0.25, *P* = 1.8 × 10^−3^, q = 0.025), *Lachnospiraceae_ND3007_group* (r = –0.23, *P* = 4.14 × 10^−3^, q = 0.038), *Lachnospira* (r = –0.20, *P* = 0.012, q = 0.065), *[Eubacterium]_eligens_group* (r = –0.19, *P* = 0.019, q = 0.085). These results suggested that the depletion of Lachnospiraceae genera might be related to the increase of blood pressure.

**FIG 4.**
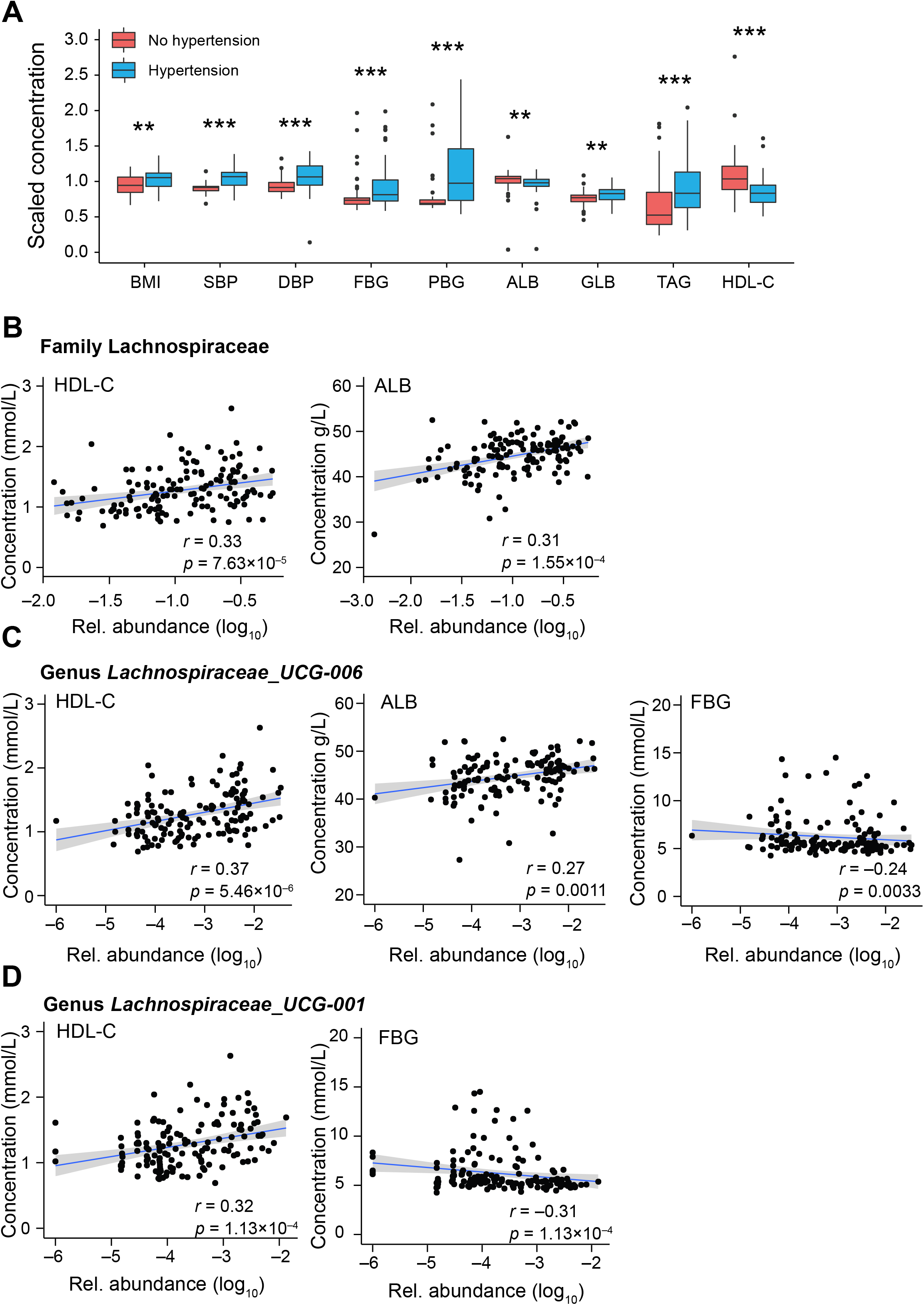
Correlations of gut microbes with hypertension-related physiological properties. (A) Boxplots show the scaled concentration of physiological properties that were significantly different between the hypertension group and the non-hypertension group (*P* < 0.05, q < 0.05, Mann-Whitney-Wilcoxon test). *, *P* < 0.05; **, *P* < 0.01, ***, *P* < 0.001. Boxes represent the interquartile range between first and third quartiles and the line inside represents the median. Whiskers denote the lowest and highest values within 1.5 × interquartile range from the first and third quartiles, respectively. (B, C, and D) Scatter plot of the concentration of log_10_-transformed relative abundance of gut microbes (X-axis) and physiological indexes (Y-axis). The blue line is plotted using linear regression, with 95% point wise confidence interval band shaded gray. The correlation and the statistical significance were calculated using Spearman correlation.

Next, we focused on the two genera of family Lachnospiraceae, including genus *Lachnospiraceae_UCG-006* and genus *Lachnospiraceae_UCG-001* whose depletion was the most prominent change in Han- and Yugur-hypertension microbiota, respectively. Family Lachnospiraceae was found positively correlated with concentration of high-density lipoprotein cholesterol (HDL-C, r = 0.33, *P* = 7.63 × 10^−5^, q = 9.30 × 10^−3^) and albumin (ALB, r = 0.31, *P* = 1.55 × 10^−4^, q = 9.30 × 10^−3^) (**Fig. 4B**). Genus *Lachnospiraceae_UCG-006* was found also positively correlated with concentration of HDL-C (r = 0.37, *P* = 5.46 × 10^−6^, q = 1.52 × 10^−3^) and ALB (r = 0.27, *P* = 1.1 × 10^−3^, q = 0.019), while negatively correlated with concentration of fasting blood glucose (FBG, r = –0.24, P = 3.3 × 10^−3^, q = 0.033) (**Fig. 4C**). Genus *Lachnospiraceae_UCG-001* was found positively correlated with concentration of HDL-C (r = 0.32, *P* = 1.13 × 10^−4^, q = 3.94 × 10^−3^) and FBG (r = –0.31, *P* = 1.13 × 10^−4^, q = 3.94 × 10^−3^). Notably, abundance of all these three microbial features were positively correlated the concentration of HDL-C. Moreover, HDL-C in our data was found significantly negatively correlated with SBP (r = –0.19, *P* = 0.023), while not significantly correlated with DBP, which was consistent with a previous study on 4552 individuals of a Korean cohort (25).

Furthermore, it has been reported that butyrate can regulate the secretion of apolipoprotein A-IV (apoA-IV), a lipid-binding protein, which modulated reverse cholesterol transport to increase serum HDL-C (26). We then randomly selected 17 individuals to test the content of serum butyrate. We found both Han and Yugur people with hypertension had lower content of butyrate than that of people without hypertension: 67.21 ± 4.23ng/ml in four Han individuals with hypertension vs. 81.66 ± 3.06ng/ml in four Han individuals without hypertension (*P* = 1.99 × 10^−3^); and 60.88 ± 8.25ng/ml in four Yugur individuals with hypertension vs. 76.21 ± 1.76ng/ml in five Yugur individuals without hypertension (*P* = 0.031).

## DISCUSSION

In this study, we found that people with hypertension and high-calorie diet exhibited gut microbial dysbiosis, represented by the considerable depletion of Lachnospiraceae genera. Moreover, we found the depletion of Lachnospiraceae were correlated with the decrease of HDL-C and increase of SBP and DBP. Furthermore, we validated that the Han people and Yugur people with hypertension had a lower serum butyrate content. We concluded that that the depletion of Lachnospiraceae genera might lead to less production of intestinal butyrate in people with hypertension, contributing to the lower level of HDL-C, elevating blood pressure and promoting hypertension (**Fig. 1C**).

A diet rich in fruits, vegetables, and low-fat dairy products and reduced saturated and total fat was recommended for people to ameliorate hypertension (8). However, the recruited cohort in our study shared a nearly opposite dietary custom with excessive intake of meat but limited intake of vegetables or fruits, which might lead to their higher incidence of hypertension. Due to a paucity of vegetables in the diet, the sufficient fermentation of plant polysaccharides was essential. Family Lachnospiraceae could ferment diverse plant polysaccharides to SCFAs (19, 21, 22, 27), which played a role in maintenance of health such as energy supply and immunity regulation (28, 29). We found that the bulk of the considerably decreased microbes in the hypertension group came from the family Lachnospiraceae, which was the most notable feature of microbial variation under the effects of hypertension related to a high-calorie diet. Nonetheless, the overall abundance of family Lachnospiraceae was not significantly decreased in Yugur-hypertension microbiota, even though it was considerably decreased in Han-hypertension microbiota. It was speculated that when hypertension developed, Han gut microbiota might be more vulnerable than Yugur gut microbiota, possibly due to the shorter time of their ancestry residence in Sunan County or the host-genetic distinction. However, this speculation was limited by the different sample size in this study, which required further investigations. Moreover, although the family Lachnospiraceae remained abundant, the genus *Lachnospiraceae UIG-001* was significantly depleted in Yugur-hypertension microbiota, but not in Han-hypertension microbiota. On the contrary, another genus, *Lachnospiraceae UIG-006*, was considerably reduced in Han-hypertension microbiota but not in Yugur-hypertension microbiota. Thus, the gut microbiota of two ethnic groups might respond differently to the stress of hypertension, which might explain the disparity in hypertension prevalence of these two ethnic groups in the community.

Intestinal butyrate, accounting for 95% of the SCFAs produced by the gut microbiota (30), was reported to increase serum HDL-C, by regulating the secretion of apoA-IV^27^. In addition, HDL-C was reported to be negatively correlated with SBP but not DBP in a cohort of 4,552 Korean (25), which was consistent with our findings. Moreover, HDL-C also played an important role in reducing the risk of a variety of cardiovascular diseases (31). Therefore, HDL-C might be a crucial link between microbes and hypertension. In this study, we proposed a potential mechanism that depletion of members from family Lachnospiraceae caused less production of intestinal butyrate in people with hypertension (19, 21, 22, 27), which might contribute to the lower level of HDL-C, elevated SBP, and hypertension development. This mechanism would be significant for people who followed a high-calorie diet, with limited intake of vegetables. Besides the link between microbes, HDL-C, and SBP, we also found a certain number of direct correlations of microbes with SBP and DBP, such as the positive correlation of SBP with *Escherichia-Shigell*. Additionally, several increased intestinal microbes such as *Sphingomonas* were also reported to be increased in blood of people with hypertension. Hence, further investigations were required for exploring multi potential mechanisms in this study, such as the microbes-metabolite (butyrate, HDL-C)-SBP-hypertension link, microbes-DBP-hypertension link, and the intestinal-blood-microbes-hypertension link.

This study also has limitations. First, since the different cohort size of Han and Yugur people could influence the statistical significance in microbial alterations, we may not be able to give a strong conclusion about the ethnic differences in microbial dysbiosis. Second, we only tested the serum butyrate content of a part of individuals in this study owing to the strict policy for blood test in Sunan County. However, these limitations would not negate the substantial microbial differentiation between people with and without hypertension under a high-calorie diet, as well as the strong correlations between the dysbiosis and HDL-C, which was of clinical importance. Moreover, the butyrate might be one of the potential media through which microbes adjusted HDL-C, and further researches into the underlying mechanisms are necessary.

### Conclusions

this study demonstrates that individuals with hypertension under a high-calorie diet exhibit a substantial depletion of Lachnospiraceae genera, which might promote hypertension progression by lowering serum HDL-C levels. This study provides a new insight into the link between microbial dysbiosis and hypertension under high-calorie diet. Further investigations on the role of gut microbiota in HDL-C regulation in a variety of cardiovascular diseases are warranted.

## MATERIALS AND METHODS

### Ethnical statement

All procedures performed in this study were approved by the Medical Ethics Committee of Northwest Minzu University (No. XBMZ-YX-202004), and in accordance with the Helsinki Declaration of 1975. All participants have provided written informed consent to take part in the study.

### Epidemiological survey

A total of 1,089 individuals aged over 18 years old from the Han (n = 639) and the Yugur (n = 450) in Sunan County were randomly selected for the epidemiological survey in 2015. Informed consent was obtained and the collection contents were as follows: 1) General information, physical health, lifestyle and behaviour, smoking history, drinking history, and family history were collected using self-designed questionnaire; 2) Physical parameters such as blood pressure, height, weight, waist circumference and body mass index (BMI) were measured using a unified method; 3) Physiological parameters such as serum total cholesterol (TC), triglyceride (TG), high density lipoprotein cholesterol (HDL-C), low density lipoprotein cholesterol (LDL-C) were detected.

### Fecal sample collection

A total of 153 fecal samples of Yugur and Han people, who had been living in Sunan County for more than 15 years, were collected in 2020. A total of 10g of feces for each sample was collected into a stool storage tube containing stool preservation fluid in the morning. The preservation fluid and stool were mixed evenly before the sample was frozen in a −80 °C freezer for ≥24 h. Within one week, we shipped samples in dry ice to the laboratory for following experiments. All of the participants must not have any taken antibiotics, microbial preparations, or antidiarrheal or weight loss drugs, and must not have a history of diarrhea or other gastrointestinal (GI) diseases, within the last month. The dietary information of these 153 individuals was collected.

### DNA extraction and 16S rRNA gene sequencing

A PowerSoil Deoxyribonucleic Acid (DNA) Extraction Kit (QIAGEN, Hilden, Germany) was used to extract genomic DNA from fecal samples according to the manufacturer’s instructions. We used 1% agarose gel electrophoresis and a NanoDrop2000 spectrophotometer (Thermo Fisher Scientific, Waltham, MA, USA) to measure DNA concentration and purity, respectively. A suitable amount of sample was added to a centrifuge tube, and sterile water was used to dilute the sample to 1 ng/ul. We used the diluted DNA as a template and specific primers 343F (5′- TACGGRAGGCAGCAG-3′) and 798R (5′-AGGGTATCTAATCCT-3′) with Tks Gflex DNA Polymerase for polymerase chain reaction (PCR) amplification of the 16S V3–V4 region in samples to ensure amplification efficiency and accuracy. The first round of PCR amplification conditions consisted of pre-denaturation at 94 °C for 5 min; followed by 26 cycles of 94 °C for 30 s, 56 °C for 30 s, and 72 °C for 20 s; and then a final extension of 72 °C for 5 min and holding at 4 °C. The second round consisted of pre-denaturation at 94 °C for 5 min; followed by seven cycles of 94 °C for 30 s, 56 °C for 30 s, and 72 °C for 20 s; and then a final extension of 72 °C for 5 min and holding at 4 °C. lllumina MiSeq sequencing (Illumina, San Diego, CA, USA) was used to generate paired-end (PE) sequences.

### Sequence process

Trimmomatic software version 0.35 was used to remove the sequences with moving windows whose mean base quality < 20 and the sequences with length < 50 bp (32). Fast Length Adjustment of SHort reads (FLASh) software version 1.2.11 was used to join PE sequences after removing impurities (33). The parameters used for joining were as follows: minimum overlap, 10 bp; maximum overlap, 200 bp; maximum rate, 20%. Quantitative Insights Into Microbial Ecology (QIIME) split_libraries.py software version 1.8.0 was used to remove PE sequences containing N bases and sequences with a base quality score Q20 < 75% (34). UCHIME software version 2.4.2 was used to remove chimeras from the remained sequences (35). Vsearch software version 2.4.2 was used for OTU clustering with 97% similarity (36), and the sequence with the greatest abundance in each OTU was taken as the representative sequence for the RDP classifier (37). A naïve Bayesian classification algorithm was used to align and annotate representative sequences. Rarefaction was set at 35,490 reads based on the curve plateaus for alpha diversity.

### Principle coordinate analysis

Unweighted and weighted Unifrac distances between OTUs among all samples were used for principal coordinate analysis (PCoA). R function “dudi.pco” in the R package “ade4” was used to perform PCoA, and R package “ggplot2” was used to visualize the results.

### Identification of the microbial biomarkers

Random forest algorithm was used to identify the microbial biomarkers for each of groups. Ten randomized 10-fold cross validation was performed on microbial features to determine the mean decrease of Gini score as the feature importance. Area Under the Receiver Operating Characteristic curve (AUROC) was calculated by performing random forest algorithm on all samples, Yugur samples, and Han samples, respectively.

### Quantitative detection of serum butyrate

80μL ice-cold acetonitrile-water (1:1, v/v, containing [^2^H9]-Pentanoic acid, [^2^H11]-Hexanoic Acid) was added into the 80mg freeze-dried serum samples. Samples were extracted by ultrasonic for 10 min in ice-water bath. Samples were then centrifuged at 4°C (12,000 rpm) for 10 min. For derivatization, 80μL of the standard solution or 80μL of the supernatants were mixed with 40μL of 200 mM 3-NPH in 50% aqueous acetonitrile and 40μL of 120 mM EDC-6% pyridine solution in the same solvent. The mixture reacted at 40℃ for 30 min. Afterward samples were placed at ice for 1 min, and then filtered through a 0.22 μm organic phase pinhole filter for subsequent UPLC-MS/MS analysis. A pooled sample for quality control was prepared by mixing aliquots of all the samples. Mixed standard stock solution was prepared and diluted to produce the calibration curve.

Liquid chromatography was performed on an Nexera UHPLC LC-30A (SHIMADZU). ACQUITY UPLC BEH C18 (100*2.1mm,1.7μm) was used for analysis. Injection volume was 2μL. The mobile phase A was water containing 0. 1% formic acid, and the mobile phase B was acetonitrile. A gradient elution procedure was used: 0 min A/B (90:10, V/V), 1 min A/B (90:10, V/V), 2min A/B (75:25, V/V), 6min A/B (65:35, V/V), 6.5 min A/B (5:95, V/V), 7.8 min A/B (5:95, V/V), 7.81 min A/B (90:10, V/V), 8.5 min A/B (90:10, V/V). All the samples were kept at 4℃ during the analysis and the column temperature was set at 40℃. Mass spectrometry was performed on the AB SCIEX Selex ION Triple Quad™ 5500 System. SCIEX OS-MQ software was used for quantification. Concentration of butyrate was calculated according to the peak area and the calibration curve.

### Statistical Analysis

For categorical metadata, samples were pooled into bins (Hypertension/Non-hypertension, Yugur people/Han people, Yugur-hypertension/Han-hypertension/Yugur-non-hypertension/Han-non-hypertension) and significant were calcultaed using Mann-Whitney-Wilcoxon Test (P values) with Benjamini and Hochberg correction (FDR, q values). Permutational multivariate analysis of variance (PERMANOVA) was performed with 9,999 permutations using unweighted-Unifrac distance matrix. Age, gender, and body mass index were used as co-variants. The significance of comparisons of serum butyrate content was calculated using *t*-test. The correlations between variables were tested using Spearman correlation.

### Data availability

Sequencing data are available in the Genome Sequence Archive (GSA) section of National Genomics Data Center (project accession number CRA005607). Data link is for review: https://ngdc.cncb.ac.cn/gsa/s/9Ja065bX.

## SUPPLEMENTARY MATERIAL

**TABLE S1** to **S9,** EXCEL file, 0.1 MB.

**TABLE S1** Epidemic survey of Hypertension prevalence and dietary structure of Han and Yugur people in Sunan County.

**TABLE S2** Permutational multivariate analysis of variance (PERMANOVA) table for 153 individuals from Han and Yugur ethnic groups.

**TABLE S3** Comparisons of taxonomic abundances between non-hypertension group and hypertension group.

**TABLE S4** Comparisons of hypertension-related genera abundances among a variety of groups.

**TABLE S5** Mean Gini importance produced by random forest algorithm on all samples.

**TABLE S6** Mean Gini importance produced by random forest algorithm on Han samples.

**TABLE S7** Mean Gini importance produced by random forest algorithm on Yugur samples.

**TABLE S8** Comparisons of physiological properties between hypertension and non-hypertension samples or between Han samples and Yugur samples.

**TABLE S9** The correlations of physiological properties with gut microbes.

## Acknowledgements

This work was partially supported by Innovation Team Training Project of Northwest Minzu University of Central Universities Basic Research Funds (Grant Nos. 31920190030), Key Project of Northwest Minzu University of Central Universities Basic Research Funds (Grant No. 31920190100), National Natural Science Foundation of China (Grant Nos. 32071465, 31871334 and 31671374), and the National Key R&D Program of China (Grant No. 2018YFC0910502).

## Author contributions

Y.L., K.N., and M.C. designed the study, conducted the data analysis, and wrote the manuscript. Y.L., K.N., M.C., Y.M., J.Z., C.C., X.Y., F.A, Z.Z., and Y.A. collected the samples, conducted the experiments, and participated in data analysis. Y.L., K.N., and M.C. supervised the study and revised the manuscript.

## References

1 Brown JM, Hazen SL. 2018. Microbial modulation of cardiovascular disease. Nat Rev Microbiol 16:171–181.

2 Yan Q, Gu Y, Li X, Yang W, Jia L, Chen C, Han X, Huang Y, Zhao L, Li P, Fang Z, Zhou J, Guan X, Ding Y, Wang S, Khan M, Xin Y, Li S, Ma Y. 2017. Alterations of the Gut Microbiome in Hypertension. Front Cell Infect Microbiol 7:381.

3 Go AS, Mozaffarian D, Roger VL, Benjamin EJ, Berry JD, Borden WB, Bravata DM, Dai S, Ford ES, Fox CS, Franco S, Fullerton HJ, Gillespie C, Hailpern SM, Heit JA, Howard VJ, Huffman MD, Kissela BM, Kittner SJ, Lackland DT, Lichtman JH, Lisabeth LD, Magid D, Marcus GM, Marelli A, Matchar DB, McGuire DK, Mohler ER, Moy CS, Mussolino ME, Nichol G, Paynter NP, Schreiner PJ, Sorlie PD, Stein J, Turan TN, Virani SS, Wong ND, Woo D, Turner MB. 2013. Heart disease and stroke statistics--2013 update: a report from the American Heart Association. Circulation 127:e6–e245.

4 Lim SS, Vos T, Flaxman AD, Danaei G, Shibuya K, Adair-Rohani H, Amann M, Anderson HR, Andrews KG, Aryee M, Atkinson C, Bacchus LJ, Bahalim AN, Balakrishnan K, Balmes J, Barker-Collo S, Baxter A, Bell ML, Blore JD, Blyth F, Bonner C, Borges G, Bourne R, Boussinesq M, Brauer M, Brooks P, Bruce NG, Brunekreef B, Bryan-Hancock C, Bucello C, Buchbinder R, Bull F, Burnett RT, Byers TE, Calabria B, Carapetis J, Carnahan E, Chafe Z, Charlson F, Chen H, Chen JS, Cheng AT, Child JC, Cohen A, Colson KE, Cowie BC, Darby S, Darling S, Davis A, Degenhardt L, et al. 2012. A comparative risk assessment of burden of disease and injury attributable to 67 risk factors and risk factor clusters in 21 regions, 1990-2010: a systematic analysis for the Global Burden of Disease Study 2010. Lancet 380:2224–60.

5 Ardissino D, Berzuini C, Merlini PA, Mannuccio Mannucci P, Surti A, Burtt N, Voight B, Tubaro M, Peyvandi F, Spreafico M, Celli P, Lina D, Notarangelo MF, Ferrario M, Fetiveau R, Casari G, Galli M, Ribichini F, Rossi ML, Bernardi F, Marziliano N, Zonzin P, Mauri F, Piazza A, Foco L, Bernardinelli L, Altshuler D, Kathiresan S. 2011. Influence of 9p21.3 genetic variants on clinical and angiographic outcomes in early-onset myocardial infarction. J Am Coll Cardiol 58:426–34.

6 Ripatti S, Tikkanen E, Orho-Melander M, Havulinna AS, Silander K, Sharma A, Guiducci C, Perola M, Jula A, Sinisalo J, Lokki ML, Nieminen MS, Melander O, Salomaa V, Peltonen L, Kathiresan S. 2010. A multilocus genetic risk score for coronary heart disease: case-control and prospective cohort analyses. Lancet 376:1393–400.

7 Yu E, Rimm E, Qi L, Rexrode K, Albert CM, Sun Q, Willett WC, Hu FB, Manson JE. 2016. Diet, Lifestyle, Biomarkers, Genetic Factors, and Risk of Cardiovascular Disease in the Nurses’ Health Studies. Am J Public Health 106:1616–23.

8 Bazzano LA, Green T, Harrison TN, Reynolds K. 2013. Dietary approaches to prevent hypertension. Curr Hypertens Rep 15:694–702.

9 Rothschild D, Weissbrod O, Barkan E, Kurilshikov A, Korem T, Zeevi D, Costea PI, Godneva A, Kalka IN, Bar N, Shilo S, Lador D, Vila AV, Zmora N, Pevsner-Fischer M, Israeli D, Kosower N, Malka G, Wolf BC, Avnit-Sagi T, Lotan-Pompan M, Weinberger A, Halpern Z, Carmi S, Fu J, Wijmenga C, Zhernakova A, Elinav E, Segal E. 2018. Environment dominates over host genetics in shaping human gut microbiota. Nature 555:210–215.

10 Wu GD, Chen J, Hoffmann C, Bittinger K, Chen YY, Keilbaugh SA, Bewtra M, Knights D, Walters WA, Knight R, Sinha R, Gilroy E, Gupta K, Baldassano R, Nessel L, Li H, Bushman FD, Lewis JD. 2011. Linking long-term dietary patterns with gut microbial enterotypes. Science 334:105–8.

11 Miyamoto J, Kasubuchi M, Nakajima A, Irie J, Itoh H, Kimura I. 2016. The role of short-chain fatty acid on blood pressure regulation. Curr Opin Nephrol Hypertens 25:379–83.

12 Wang Z, Chen Z, Zhang L, Wang X, Hao G, Zhang Z, Shao L, Tian Y, Dong Y, Zheng C, Wang J, Zhu M, Weintraub WS, Gao R. 2018. Status of Hypertension in China: Results From the China Hypertension Survey, 2012-2015. Circulation 137:2344–2356.

13 Society CN. 2008. The Pagoda of Balanced Diet for Chinese Residents. Acta Pharmacol Sin 30(1): 14–15.

14 Louca P, Nogal A, Wells PM, Asnicar F, Wolf J, Steves CJ, Spector TD, Segata N, Berry SE, Valdes AM, Menni C. 2021. Gut microbiome diversity and composition is associated with hypertension in women. J Hypertens 39:1810– 1816.

15 Clark WF, Sontrop JM, Macnab JJ, Salvadori M, Moist L, Suri R, Garg AX. 2010. Long term risk for hypertension, renal impairment, and cardiovascular disease after gastroenteritis from drinking water contaminated with Escherichia coli O157:H7: a prospective cohort study. BMJ 341:c6020.

16 Schierwagen R, Alvarez-Silva C, Madsen MSA, Kolbe CC, Meyer C, Thomas D, Uschner FE, Magdaleno F, Jansen C, Pohlmann A, Praktiknjo M, Hischebeth GT, Molitor E, Latz E, Lelouvier B, Trebicka J, Arumugam M. 2019. Circulating microbiome in blood of different circulatory compartments. Gut 68:578–580.

17 Jing Y, Zhou H, Lu H, Chen X, Zhou L, Zhang J, Wu J, Dong C. 2021. Associations Between Peripheral Blood Microbiome and the Risk of Hypertension. Am J Hypertens 34:1064–1070.

18 Santisteban MM, Qi Y, Zubcevic J, Kim S, Yang T, Shenoy V, Cole-Jeffrey CT, Lobaton GO, Stewart DC, Rubiano A, Simmons CS, Garcia-Pereira F, Johnson RD, Pepine CJ, Raizada MK. 2017. Hypertension-Linked Pathophysiological Alterations in the Gut. Circ Res 120:312–323.

19 Byndloss MX, Olsan EE, Rivera-Chávez F, Tiffany CR, Cevallos SA, Lokken KL, Torres TP, Byndloss AJ, Faber F, Gao Y, Litvak Y, Lopez CA, Xu G, Napoli E, Giulivi C, Tsolis RM, Revzin A, Lebrilla CB, Bäumler AJ. 2017. Microbiota-activated PPAR-γ signaling inhibits dysbiotic Enterobacteriaceae expansion. Science 357:570–575.

20 Buffie CG, Bucci V, Stein RR, McKenney PT, Ling L, Gobourne A, No D, Liu H, Kinnebrew M, Viale A, Littmann E, van den Brink MR, Jenq RR, Taur Y, Sander C, Cross JR, Toussaint NC, Xavier JB, Pamer EG. 2015. Precision microbiome reconstitution restores bile acid mediated resistance to Clostridium difficile. Nature 517:205–8.

21 Rivera-Chávez F, Zhang LF, Faber F, Lopez CA, Byndloss MX, Olsan EE, Xu G, Velazquez EM, Lebrilla CB, Winter SE, Bäumler AJ. 2016. Depletion of Butyrate-Producing Clostridia from the Gut Microbiota Drives an Aerobic Luminal Expansion of Salmonella. Cell Host Microbe 19:443–54.

22 Sorbara MT, Littmann ER, Fontana E, Moody TU, Kohout CE, Gjonbalaj M, Eaton V, Seok R, Leiner IM, Pamer EG. 2020. Functional and Genomic Variation between Human-Derived Isolates of Lachnospiraceae Reveals Inter- and Intra-Species Diversity. Cell Host Microbe 28:134–146.e4.

23 Studer N, Desharnais L, Beutler M, Brugiroux S, Terrazos MA, Menin L, Schürch CM, McCoy KD, Kuehne SA, Minton NP, Stecher B, Bernier-Latmani R, Hapfelmeier S. 2016. Functional Intestinal Bile Acid 7α-Dehydroxylation by Clostridium scindens Associated with Protection from Clostridium difficile Infection in a Gnotobiotic Mouse Model. Front Cell Infect Microbiol 6:191.

24 Kasahara K, Krautkramer KA, Org E, Romano KA, Kerby RL, Vivas EI, Mehrabian M, Denu JM, Bäckhed F, Lusis AJ, Rey FE. 2018. Interactions between Roseburia intestinalis and diet modulate atherogenesis in a murine model. Nat Microbiol 3:1461–1471.

25 Cho KH, Park HJ, Kim JR. 2020. Decrease in Serum HDL-C Level Is Associated with Elevation of Blood Pressure: Correlation Analysis from the Korean National Health and Nutrition Examination Survey 2017. Int J Environ Res Public Health 17:1101.

26 Nazih H, Nazih-Sanderson F, Krempf M, Michel Huvelin J, Mercier S, Marie Bard J. 2001. Butyrate stimulates ApoA-IV-containing lipoprotein secretion in differentiated Caco-2 cells: role in cholesterol efflux. J Cell Biochem 83:230–8.

27 Boutard M, Cerisy T, Nogue PY, Alberti A, Weissenbach J, Salanoubat M, Tolonen AC. 2014. Functional diversity of carbohydrate-active enzymes enabling a bacterium to ferment plant biomass. PLoS Genet 10:e1004773.

28 Arpaia N, Campbell C, Fan X, Dikiy S, van der Veeken J, deRoos P, Liu H, Cross JR, Pfeffer K, Coffer PJ, Rudensky AY. 2013. Metabolites produced by commensal bacteria promote peripheral regulatory T-cell generation. Nature 504:451–5.

29 Pascale A, Marchesi N, Marelli C, Coppola A, Luzi L, Govoni S, Giustina A, Gazzaruso C. 2018. Microbiota and metabolic diseases. Endocrine 61:357–371.

30 He J, Zhang P, Shen L, Niu L, Tan Y, Chen L, Zhao Y, Bai L, Hao X, Li X, Zhang S, Zhu L. 2020. Short-Chain Fatty Acids and Their Association with Signalling Pathways in Inflammation, Glucose and Lipid Metabolism. Int J Mol Sci 21:6356.

31 Mahdy Ali K, Wonnerth A, Huber K, Wojta J. 2012. Cardiovascular disease risk reduction by raising HDL cholesterol--current therapies and future opportunities. Br J Pharmacol 167:1177–94.

32 Bolger AM, Lohse M, Usadel B. 2014. Trimmomatic: a flexible trimmer for Illumina sequence data. Bioinformatics 30:2114–20.

33 Magoč T, Salzberg SL. 2011. FLASH: fast length adjustment of short reads to improve genome assemblies. Bioinformatics 27:2957–63.

34 Caporaso JG, Kuczynski J, Stombaugh J, Bittinger K, Bushman FD, Costello EK, Fierer N, Pena AG, Goodrich JK, Gordon JI, Huttley GA, Kelley ST, Knights D, Koenig JE, Ley RE, Lozupone CA, McDonald D, Muegge BD, Pirrung M, Reeder J, Sevinsky JR, Turnbaugh PJ, Walters WA, Widmann J, Yatsunenko T, Zaneveld J, Knight R. 2010. QIIME allows analysis of high-throughput community sequencing data. Nat Methods 7:335–6.

35 Edgar RC, Haas BJ, Clemente JC, Quince C, Knight R. 2011. UCHIME improves sensitivity and speed of chimera detection. Bioinformatics 27:2194– 2200.

36 Rognes T, Flouri T, Nichols B, Quince C, Mahé F. 2016. VSEARCH: a versatile open source tool for metagenomics. PeerJ 4:e2584.

37 Wang Q, Garrity GM, Tiedje JM, Cole JR. 2007. Naive Bayesian classifier for rapid assignment of rRNA sequences into the new bacterial taxonomy. Appl Environ Microbiol 73:5261–5267.

